# Tau-induced ribosomal collisions impair memory through the activation of the integrated stress response

**DOI:** 10.64898/2026.01.12.699151

**Authors:** Harrison T. Evans, Srinidhi V. Kalavai, Victor Wu, Ambika Polavarapu, Emma Balamoti, Wendy J Liu, Olivia Mosto, Drew Alder, Vasvi Jain, Eléa Denil, Astra Yu, Esther Assenso, Mauricio Oliveria, Taylor Moniz, Ela N. Golhan, Xunzhao Zhang, Charles J. Sheehan, Eric Klann

**Author notes:** Co-first authors (Harrison T. Evans, Srinidhi V. Kalavai).

## Abstract

The formation of new long-term memories is reliant upon the spatial and temporal regulation of mRNA translation. Translational control has been demonstrated to be disrupted in neurodegenerative diseases, which exhibit impairments in both homeostatic translation and memory formation, such as Alzheimer’s disease (AD) and frontotemporal dementia (FTD). However, the precise mechanisms by which this dysregulation occurs, as well as the pathogenic consequences of this dysregulation have yet to be described. Here we establish that FTD-associated tau mutations impair protein synthesis prior to the onset of memory impairments by slowing ribosomal elongation speed, causing ribosomes to collide upon mRNAs. We reveal that this tau-induced ribosomal collision ultimately impairs memory-associated translation through activation of the integrated stress response (ISR) via GCN2. Pharmacological prevention of this ISR activation not only rescues memory formation in the PS19 mouse model of FTD, but also attenuates neuronal death, decreases tau phosphorylation and accumulation, and improves survival. Collectively, our data elucidates a novel mechanism by which mRNA translation is impaired early in neurodegeneration, identifies several pathological phenotypes which are traceable to impairments in mRNA translation, and highlights the therapeutic potential of rescuing these translational impairments.

## Introduction

To facilitate the plasticity required for the formation, storage, and recall of long-term memories, the cells of the central nervous system must spatially and temporally regulate their proteomes. One of the fundamental pillars of this regulation is the translation of mRNAs into new proteins by ribosomes, a process which is metabolically demanding and as such is tightly controlled (Kapur *et al*, 2017). Indeed, brain cells such as neurons, astrocytes, and microglia exhibit a remarkable degree of translation regulation, whether it be the differential expression and localization of RNA-binding proteins (Thelen & Kye, 2020), the preference for dendritic mRNAs to undergo translation via monosomes (singular ribosomes attached to a mRNA) (Biever *et al*, 2020), or even the possibility of heterogeneous ribosomal populations (Rothschild *et al*, 2024; Fusco *et al*, 2021).

It is little wonder then that the dynamic regulation of mRNA translation plays a pivotal role in the functions of the nervous system, most notably memory. In fact, the synthesis of new proteins is so integral to the formation of new memories, that reliance upon protein synthesis is used to molecularly distinguish long-term memory from short-term memory (Davis & Squire, 1984; Shrestha *et al*, 2020a). Blocking protein synthesis prevents long-term memory formation, with both pharmacological and chemogenic methods of inhibiting protein synthesis being used to identify the specific brain regions and cell-types involved in different forms of memory (Dubue *et al*, 2015; Shrestha *et al*, 2020b).

Several pathways which regulate mRNA translation are also implicated in memory, such as the mammalian target of rapamycin complex 1 (mTORC1). During memory formation, mTORC1 activation results in increased phosphorylation of ribosomal protein s6 (RPS6) and 4E-binding proteins (Hoeffer & Klann, 2010), which enhances the translation of select mRNAs, several of which encode ribosomal proteins (RPs) and translation elongation factors. This increase in the abundance of translational machinery may then further enable the synthesis of plasticity related proteins such as BDNF (Moy *et al*, 2018) and the AMPA receptor subunits, GluA1 and GluA2 (Fortin *et al*, 2012).

Another translation regulatory pathway that has been demonstrated to be involved in memory is the integrated stress response (ISR). Canonically, the ISR is thought to act as a response to cellular stress, with four kinases, PERK, GCN2, PKR, & HRI acting as sensors for ER stress, ribosomal collision, dsRNA exposure, and heme deficiency respectively (Costa-Mattioli & Walter, 2020; Zhou *et al*, 2025) (Fig 3A). Under these conditions, these kinases increase the phosphorylation of the alpha subunit of the eukaryotic initiation factor 2 (eIF2α), which inhibits translation of mRNA’s by preventing recruitment of methionine initiator tRNA to the 40S ribosomal subunit. This reduction in general protein synthesis is paired with an increase in the translation of mRNAs that are components of rescue pathways in an attempt to resolve the cellular stressor (Shrestha & Klann, 2022). However, even under homeostatic conditions, a subset of eIF2α remains phosphorylated, with a recent study postulating that following neuronal activation, this pool of p-eIF2α is rapidly dephosphorylated by the phosphatase GADD34, potentially enabling the increase in mRNA translation that is required during the initial stages of memory consolidation (Oliveira *et al*, 2024).

Emerging evidence suggests that the regulation of mRNA translation is perturbed in several neurodegenerative diseases in which long-term memory is impaired, such as Alzheimer’s disease (AD) and frontotemporal dementia (FTD) (Storkebaum *et al*, 2023). Indeed, several aggregation prone proteins such as amyloid-β (Elder *et al*, 2021; Lourenco *et al*, 2013; Caccamo *et al*, 2010), alpha-synuclein (Khan *et al*, 2023), TDP-43 (Pisciottani *et al*, 2023), and FUS (Sévigny *et al*, 2020) have been shown to dysregulate mRNA translation, potentially through pathways such as the ISR. In addition, proteomic studies of patient brain samples have shown that several components of cellular translational machinery, such as ribosomal proteins, are depleted in early stages of these neurodegenerative diseases (Ding *et al*, 2005).

One pathogenic molecule which may impair mRNA translation is the microtubule-associated protein tau (Koren *et al*, 2020). Encoded by the *MAPT* gene, the canonical role of tau is as a regulator of microtubule stability (Wang & Mandelkow, 2016). In neurodegenerative diseases such as AD, FTD, progressive supranuclear palsy (PSP), and corticobasal degeneration (CBD), tau has been observed to fall off of microtubules, become hyperphosphorylated, and form various aggregated species (Creekmore *et al*, 2024) . In these diseases, collectively termed tauopathies, aberrant tau is thought to cause pathology by impairing several essential neuronal functions that ultimately results in cell death (Götz *et al*, 2019). Several mutations in *MAPT* have been associated with familial forms of FTD, such as P301S and V337M (Strang *et al*, 2019). Expression of this FTD-mutant tau is able to recapitulate many of the neurodegenerative phenotypes observed in patients, in model systems including *Caenorhabditis elegans* (Natale *et al*, 2020)*, Drosophila Melanogaster* (Gistelinck *et al*, 2012)*, Mus Musculus*(Götz *et al*, 2018), as well as neurons(Sohn *et al*, 2019) and cerebral organoids (Bowles *et al*, 2021) grown from patient-derived inducible pluripotent stem cells.

Emerging evidence has demonstrated that pathogenic tau dysregulates mRNA. Expression of FTD-associated mutant tau has been shown to suppress global protein synthesis and decrease both the synthesis and abundance of select ribosomal proteins (Evans *et al*, 2019; Koren *et al*, 2019). Mutant tau may also interfere with the biogenesis or recycling of ribosomes (Evans *et al*, 2021b), sequester various initiation and elongation factors(Kauwe *et al*, 2025), and alter the availability of certain mRNAs through its interaction with various RNA-binding proteins (RBPs) (Apicco *et al*, 2018; Vanderweyde *et al*, 2016).

Despite this progress, three distinct questions regarding tau-induced impairments in translation remain unanswered. First, despite numerous *de novo* proteomic studies identifying the proteins that exhibit impaired synthesis in the presence of pathogenic tau, the mechanism by which FTD-mutant tau dysregulates mRNA translation remains unclear. Second, it is unclear when in disease progression these impairments in translation are manifested and how these impairments change over time. Finally, it is unknown if these observed impairments in mRNA translation are merely a consequence of other pathogenic effects of tau, or whether tau-induced translational dysregulation causes or exacerbates aspects of tau pathology, including memory impairments.

In order to answer these questions, here we leverage several advanced translatomic techniques to characterize tau-induced translational impairments in the widely utilized PS19 mouse model of tauopathies. We reveal that prior to the onset of memory impairments, P301S-mutant tau slows the speed at which ribosomes translate mRNAs in the hippocampus of PS19 mice. We also demonstrate that FTD-mutant tau can cause ribosomes to collide upon mRNAs, the first time this phenomenon has been observed in mice or as the result of pathology. These ribosome collisions result in activation of the integrated stress response by GCN2, leading to a further decrease in global protein synthesis levels. We reveal that it is this further impairment in mRNA translation which causes memory impairment in these mice, as preventing ISR activation via GCN2 rescues memory formation in the PS19 mice. We also demonstrate that repeated inhibition of the ISR is able to alleviate other elements of tau pathology, including tau accumulation and phosphorylation, hippocampal neuronal degeneration, and survival. Collectively, our findings highlight a novel mechanism by which proteostatic and metabolic stressors can drive pathology in neurodegenerative diseases.

## Results

### Memory-associated translation is impaired in 9-month-old PS19 mice

Given the critical role of mRNA translation in facilitating the initial stages of long-term memory formation, we sought to determine if FTD-mutant tau impairs memory-associated translation. To achieve this, we elected to utilize the widely studied PS19 mouse model of FTD (Götz *et al*, 2018). These mice express the 1N4R isoform of human tau (hTau) harboring the FTD-associated mutation, P301S, under the mouse prion protein promoter (Yoshiyama *et al*, 2007). In these mice, FTD-tau becomes hyperphosphorylated and forms aggregates, leading to the development of neurofibrillary tangle (NFT)-like inclusions. As a result, these mice recapitulate many of the symptoms of FTD, including synaptic loss, gliosis, impairments in long-term potentiation, and neuronal loss (Yoshiyama *et al*, 2007; Lasagna-Reeves *et al*, 2016).

PS19 mice also exhibit cognitive impairment, showing a reduced ability to form associative memories in tasks such as contextual fear conditioning (CFC) (Lasagna-Reeves *et al*, 2016). In contextual fear conditioning, mice are trained to associate a context (i.e. being placed in a box) with receiving a mild foot shock. The ability of these mice to form and recall this associative memory is then tested by measuring the conditioned response, i.e. freezing, in a probe trial 24 hours later where mice are re-exposed to the context without receiving a shock (Fig 1A). We demonstrate that despite learning to associate the context with the shock at a similar rate to their WT littermates during training, 9-month-old PS19 mice on the C57BL/6J background exhibit reduced freezing in the shock context 24 hours later, demonstrating an impairment in the formation of long-term fear associative memory. This impairment is less pronounced at 6 months of age, only trending towards significance, and is entirely absent in 3-month-old PS19 mice (Supplementary Fig 1).

**Figure 1:**
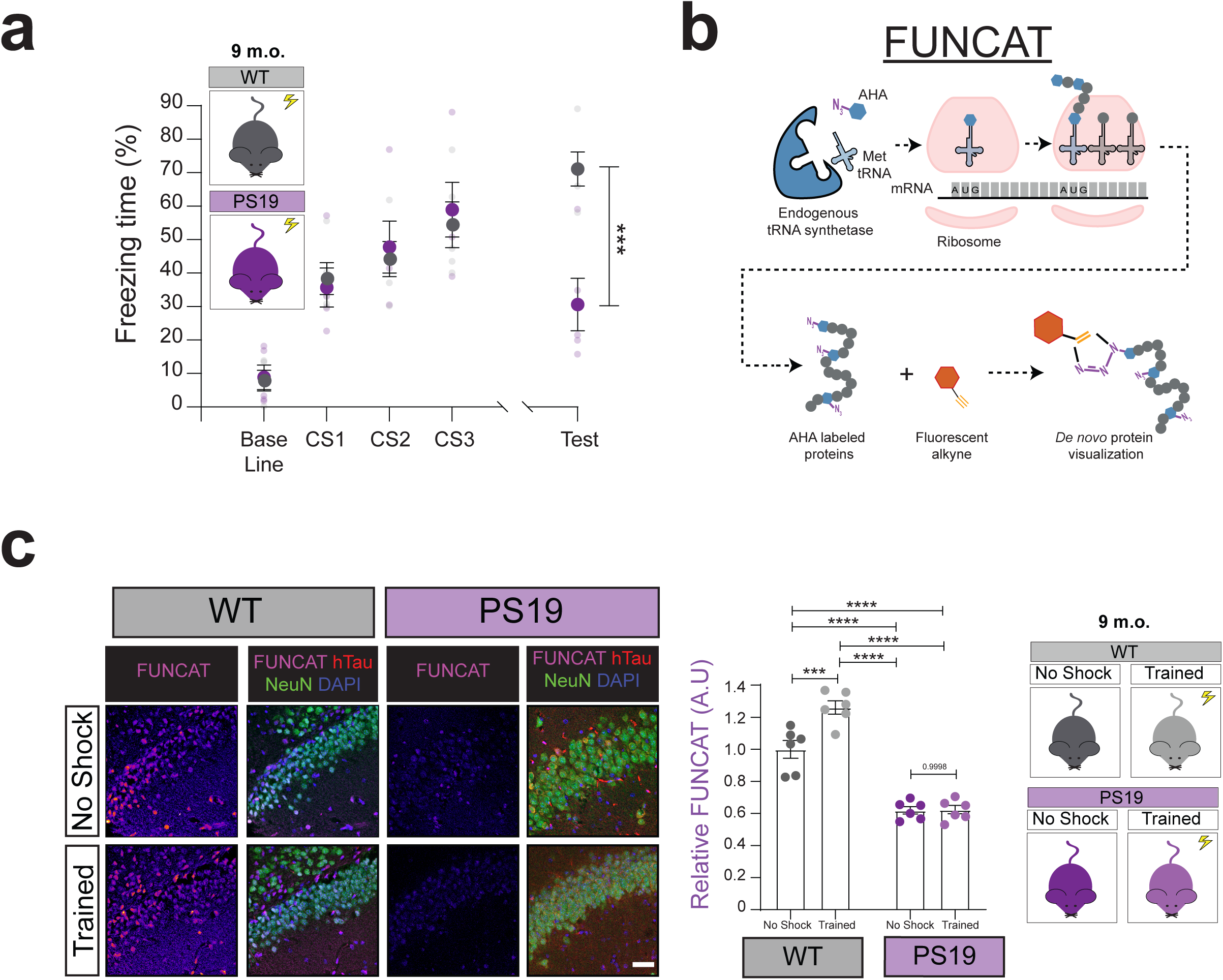
FTD-mutant tau impairs memory-associated mRNA translation in symptomatic PS19 mice. (A) 9-month-old PS19 mice exhibit memory impairments in contextual fear conditioning as shown by reduced freezing compared to WT controls in a probe test 24 hours following training (Two-way ANOVA, Sidak’s MCT n= 5 animals,***= p≤0.001). (B) Retro-orbital delivery of AHA enables the rapid visualization of the de novo proteome throughout the rodent brain. (C) 9-month-old PS19 mice exhibit impairments in memory-induced translation evidenced by the lack of increased neuronal FUNCAT signal in the hippocampus 2 hours following training via CFC (Two-way ANOVA, Sidak’s MCT n= 6 animals,***= p≤0.001, ****= p≤0.0001). Scale bar = 40 μm.

Given the impairments in long-term memory we observed in 9-month-old PS19 mice, we next sought to determine whether memory-associated mRNA translation was impaired in these mice. We recently developed a novel technique for rapidly mapping stimulus-induced changes in mRNA translation throughout the rodent brain (Evans *et al*, 2025). Here, newly synthesized proteins are labeled with the methionine surrogate, azidohomoalanine (AHA), before being visualized via fluorescence non-canonical amino acid tagging (FUNCAT) (Evans *et al*, 2021a) (Fig 1B). AHA can be delivered to awake, behaving mice via retro-orbital injection, allowing for the visualization of the *de novo* proteome in time periods as short as 30 minutes. In a previous study, this advancement allowed us to detect robust increases in protein synthesis in the 2 hours immediately following training in CFC, especially in brain regions associated with spatial memory, such as the hippocampus (Evans *et al*, 2025).

We leveraged this technical advancement to determine whether CFC-induced mRNA translation in the hippocampus is impaired in 9-month-old PS19 mice. We found that in mice not exposed to the unconditioned stimulus (i.e. no-shock mice), baseline hippocampal protein synthesis is decreased by ≈40% in PS19 mice compared to WT controls. Interestingly, unlike their WT counterparts, PS19 mice did not show an increase in hippocampal protein synthesis in the two hours following CFC training, which is indicative of an impairment in memory-associated translation (Fig 1C).

### PS19 mice exhibit impaired translation, slowed ribosomal elongation speed and collided ribosomes prior to onset of memory impairments

To elucidate if the observed impairments in memory-associated translation caused memory impairments in PS19 mice, we first sought to determine if homeostatic impairments in mRNA translation predate the development of memory impairments in these mice. We therefore used AHA labelling and FUNCAT to quantify hippocampal neuronal protein synthesis levels in 3-month-old PS19 mice, which do not exhibit impaired memory at this stage of disease progression (Supplementary Fig 1 A). Interestingly, this revealed that homeostatic mRNA translation is dysregulated prior to the onset of memory impairments in this model of FTD, with baseline protein synthesis being decreased by ≈20% in hippocampal neurons of 3-month-old PS19 mice compared to WT controls, with this decrease correlating with tau levels (Fig 2A&B).

**Figure 2:**
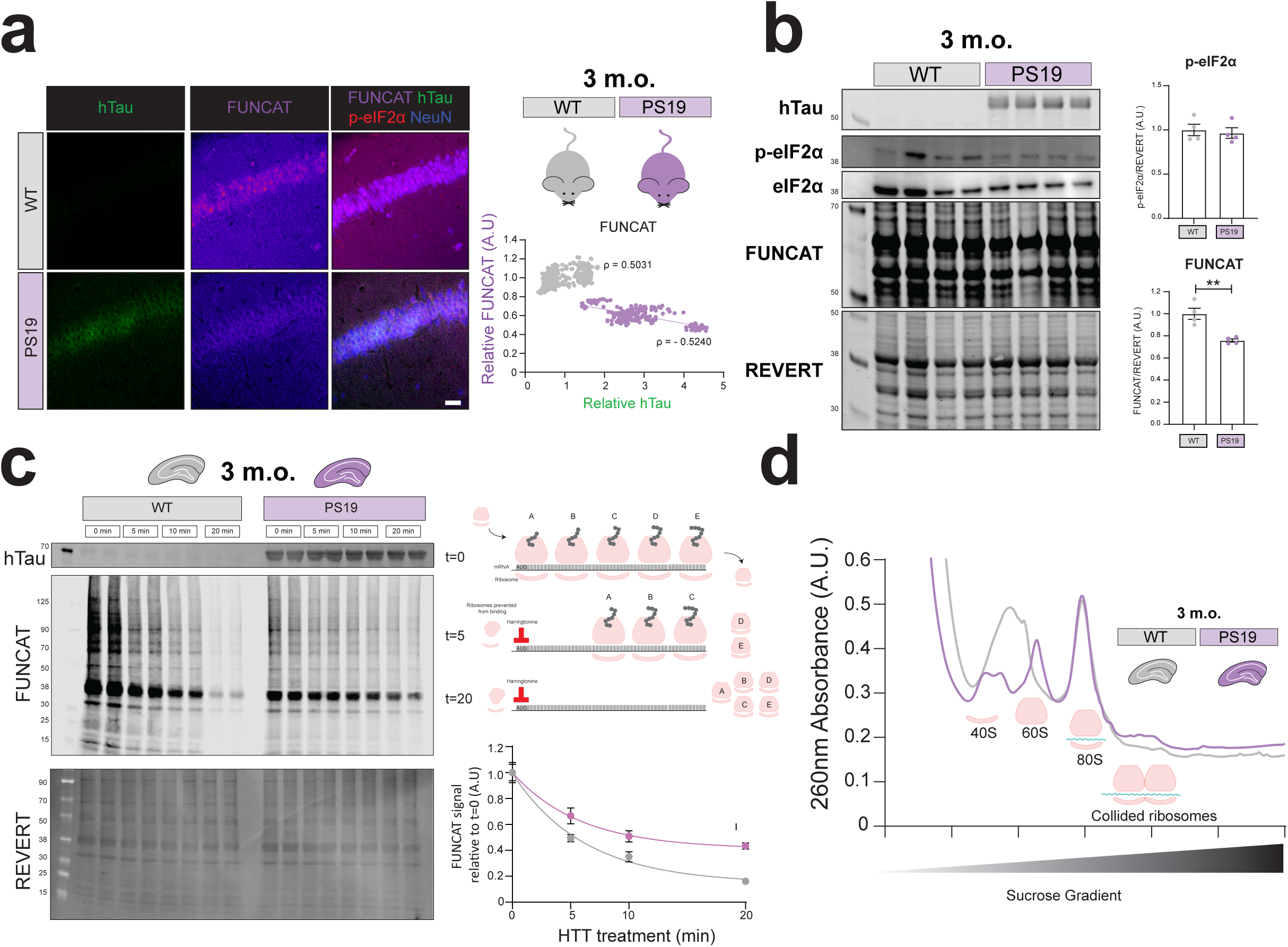
PS19 mice exhibit impaired translation, slowed ribosomal elongation speed and collided ribosomes prior to onset of memory impairments. (A) 3-month-old PS19 mice exhibit decreased homeostatic neuronal mRNA translation in the hippocampus compared to WT controls, with this decrease being correlated with human tau levels (Spearman’s correlation, n=6 animals, data points = neurons). Scale bar = 40 μm. (B) Western blot of hippocampal lysate reveals that eIF2α phosphorylation levels remain unchanged in 3-month-old PS19 mice despite the decrease in protein synthesis as measured by FUNCAT (Welch’s T.test, n=4 mice, ** = p≤0.01) (C) Harringtonine (HT) freezes ribosomes at the initiation site, preventing newly bound ribosomes from starting elongation. As a result, FUNCAT signal decreases over time as ribosomes already undergoing elongation prior to HT treatment run off the mRNA. Hippocampal slices taken from 3-month-old PS19 mice exhibit a reduced decay in FUNCAT signal (normalized to t=0), revealing a slowing of ribosomal elongation speed compared to WT controls (Single-phase logarithmic decay, ∑ of squares F-test, n=4 mice, error bars = SEM). (D) Polysome profile of RNAse digested hippocampal lysate reveals that P301S-hTau induces ribosomal collision in 3-month-old PS19 mice (n=3 mice). In polysome profiling, ribosomal subunits, monosomes and polysomes are separated on a sucrose gradient before being detected via 260nm absorbance. RNAse is used to digest the mRNA between ribosomes, resulting in the elimination of polysomes. As steric hinderance prevents RNAses from digesting mRNA between collided ribosomes, the presence of a disome peak following RNAse digestion is indicative of ribosomal collision.

We next sought to determine the mechanism by which basal mRNA translation was impaired in these mice by examining the different translational control pathways. We first examined translation initiation, i.e. the recruitment of ribosomes to the start codon. Initiation can be regulated by the phosphorylation of eIF2α through the ISR (Shrestha & Klann, 2022). Surprisingly, 3-month-old PS19 mice did not exhibit elevated levels of p-eIF2α compared to WT controls (Fig 2B, Supplementary Fig 2A). Given the sensitivity of translation initiation to even minor increases in p-eIF2α, we sought to confirm whether inhibiting the ISR rescued translation at this time point. We therefore treated 3-month-old PS19 mice with the ISR inhibitor, ISRIB (Sidrauski *et al*, 2015). ISRIB treatment was unable to rescue basal mRNA translation levels in these mice, confirming that the ISR is not activated at this stage of disease progression (Supplementary Fig 2B).

We next sought to examine the next step of mRNA translation, ribosomal elongation, using harringtonine (HT) run-off assays. HT inhibits the formation of the first peptidyl bond, preventing ribosomes from translocating from the start codon (Fresno *et al*, 1977). As such, in HT run-off assays, the only proteins which can be synthesized come from ribosomes which were already elongating prior to the addition of HT (Mohamed & Klann, 2023). By using FUNCAT to label proteins synthesized by these already elongating ribosomes and measuring how this signal decays with increasing periods of HT treatment, it is possible to indirectly measure ribosomal elongation rates, with a reduction in the decay of FUNCAT signal being indictive of slowed ribosomal elongation. Performing HT run-off assays on hippocampal slices taken from 3-month-old PS19 mice revealed that ribosomal elongation speed is slowed compared to WT controls (Fig 2C). We also showed that both P301S and V337M mutant human tau slows ribosomal elongation in transfected HEK293 cells (Supplementary Fig 3A).

Given our observation that ribosomal elongation speed is slowed in PS19 mice, we next sought to determine if these mice exhibit the recently described phenomenon of ribosomal collision. When ribosomes elongate at different rates on a mRNA strand, they can collide, activating various cellular stress pathways (Wu *et al*, 2020). To detect ribosomal collision in 3-month-old PS19 mice, we utilized polysome profiling following RNAse digestion. Here, ribosomes are separated based on size along a sucrose gradient and detected using their absorbance at 260nm. As polysomes (multiple ribosomes attached to a mRNA) are heavier than monosomes (singular ribosomes attached to an mRNA), they appear as distinct peaks further down the gradient. When RNAses are added prior to polysome profiling, the mRNA strands connecting polysomes are digested, resulting in polysomes appearing in the monosome fraction. Previous studies have established that RNAses are unable to digest the mRNA between collided ribosomes, leading to the presence of a distinct disome peak (Wolin & Walter, 1988). By performing polysome profiling on hippocampal lysate, we reveal that unlike their WT littermates, 3-month-old PS19 mice exhibit a strong disome peak after RNAse digestion, suggesting that these mice exhibit increased ribosome collisions (Fig 2D). In addition, we found that P301S-hTau increases ribosomal collision in HEK293 cells compared to non-mutant tau and a fluorescent protein control (Supplementary Fig 3B).

### Ribosomal collision induces ISR activity in PS19 mice via activation of GCN2

Previous work in cell lines has established that the collision of ribosomes can activate the ISR via increasing GCN2 activity (Zhou *et al*, 2025). To determine if this occurs in tauopathies, we examined the levels of GCN2 phosphorylation in 6-month-old PS19 mice, where ribosomal collisions appear more pronounced (Supplementary Fig 4). Western blot analysis confirmed that both GCN2 and eIF2α phosphorylation is increased in the hippocampus of 6-month-old PS19 mice compared to WT controls, suggesting that the ISR is activated by GCN2 at this stage of disease progression (Fig 3B).

**Figure 3:**
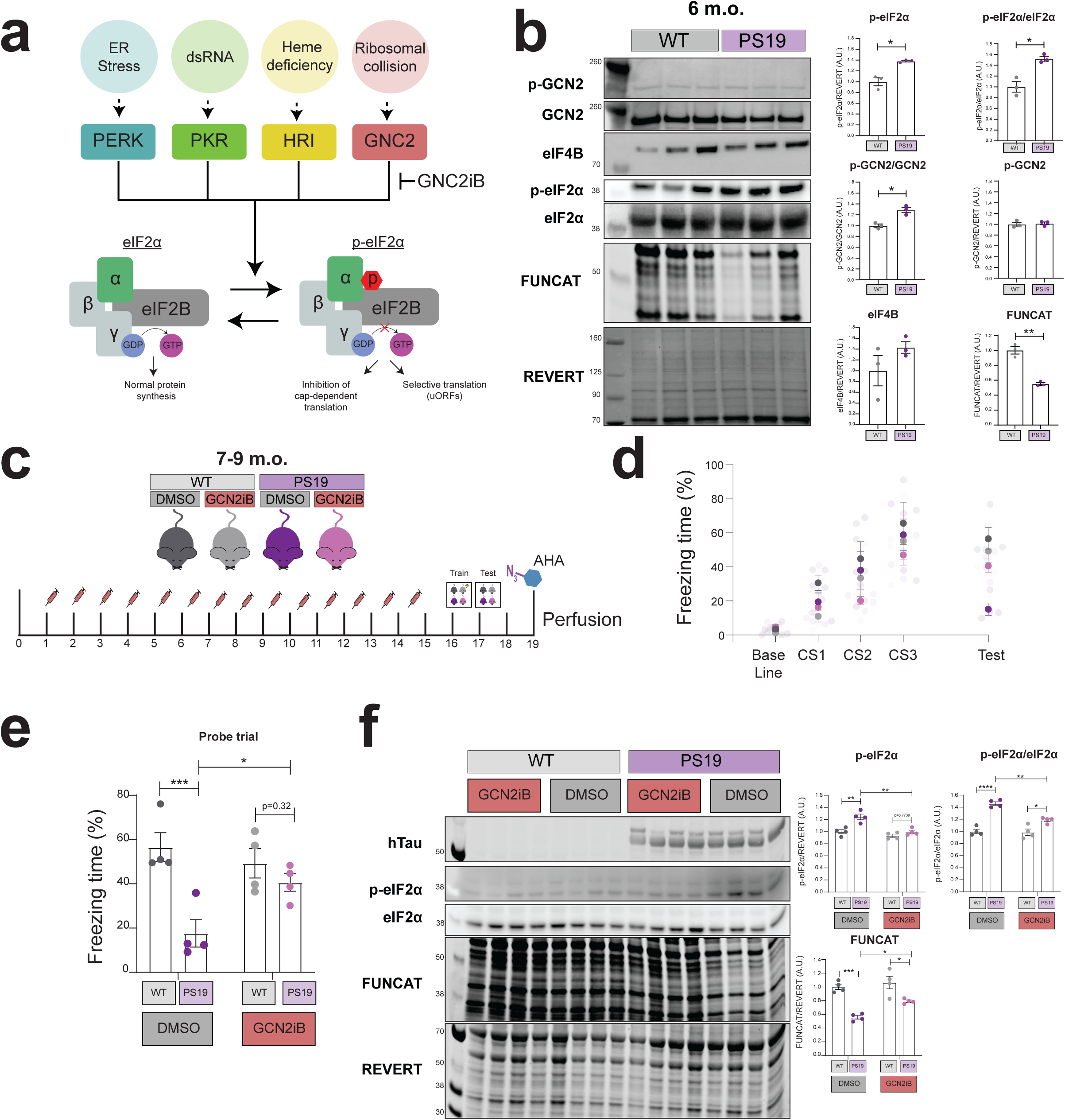
GCN2-mediated activation of the integrated stress response results in further impairments in mRNA translation in symptomatic PS19 mice, leading to impaired memory. (A) Ribosomal collision activates GCN2, leading to the phosphorylation of eIF2α and activation of the integrated stress response, which suppresses global translation and promotes cellular rescue pathways. (B) 6-month-old PS19 mice exhibit increased phosphorylation of both GCN2 and p-eIF2a (Welch’s T.test, n=3 mice *=p≤0.05,**=p≤0.01). (C) Schematic of GCN2iB treatment in 7-9-month-old PS19 mice. (D&E) Inhibition of GCN2 using GCN2iB rescues long-term memory formation following contextual fear conditioning in 7-9-month-old PS19 mice (Two-way ANOVA, Tukey’s MCT, *=p≤0.05, ***=p≤0.001, n=4 animals). (F) Pharmacological inhibition of GCN2 decreases the phosphorylation of eIF2α and partially rescues homeostatic mRNA translation in 7-9-month-old PS19 mice (Welch’s T.test, n=4 mice, *=p≤0.05,**=p≤0.01, ***=p≤0.001, ****=p≤0.0001).

We next determined whether preventing the activation of the ISR by GCN2 could rescue mRNA translation and memory. 7-9 month-old PS19 mice were treated for two weeks with the GCN2 inhibitor, GCN2iB (Nakamura *et al*, 2018), prior to contextual fear conditioning. Unlike DMSO treated controls, PS19 mice treated with GCN2iB did not show reduced freezing compared to WT mice in the probe trial (Fig 3D&E), suggesting that GCN2-mediated activation of the ISR is responsible for the memory impairments observed in PS19 mice. Biochemical analysis of these mice confirmed that inhibition of GCN2 attenuated the increase in p-eIF2α levels observed in the hippocampus of PS19 mice at this age (Fig 3F). Homeostatic mRNA translation was also partially rescued in these mice, with GCN2iB treated PS19 mice only exhibiting a ≈20% reduction in FUNCAT signal, compared to the ≈40% reduction observed in DMSO treated controls (Fig 3F). Taken together, these data suggest that GCN2-mediated activation of the ISR further inhibits mRNA translation in PS19 mice, and it is this further inhibition which ultimately causes the memory impairments observed in these mice.

### ISR inhibition attenuates multiple components of tau pathology

Given our observation that GCN2-mediated activation of the ISR causes long-term memory to be impaired in PS19 mice, we next sought to determine if other aspects of tau pathology are exacerbated by ISR activation. To achieve this, we again utilized the ISR inhibitor ISRIB, this time treating 9-month-old PS19 mice daily for two weeks, with these treatment paradigm previously being established to reduce ISR activity in AD mouse models (Oliveira *et al*, 2021). ISRIB prevents the effects of the ISR by binding to eIF2B and stabilizing it in a conformation which allows eIF2B to continue acting as a guanidine exchange factor for eIF2a leading to the recruitment of the initiator methionine tRNA to the 40S ribosomal subunit, essentially bypassing the effect of p-eIF2α (Zyryanova *et al*, 2021). We observed that in 9-month-old PS19 mice, total levels of p-eIF2α are increased compared to WT controls, with this increase being correlated with the levels of P301S-hTau and decreased mRNA translation (Fig 4A). We found that, similar to GCN2 inhibition, inhibiting the ISR partially rescues baseline hippocampal protein synthesis in 9-month-old PS19 mice, boosting FUNCAT levels to ≈80% of WT controls (Fig 4B). ISR inhibition also partially rescued long-term memory, with ISRIB-treated PS19 mice exhibiting only a ≈20% reduction in freezing compared to WT controls, whereas DMSO treated PS19 controls exhibited a ≈50% reduction in freezing (Fig 4C).

**Figure 4:**
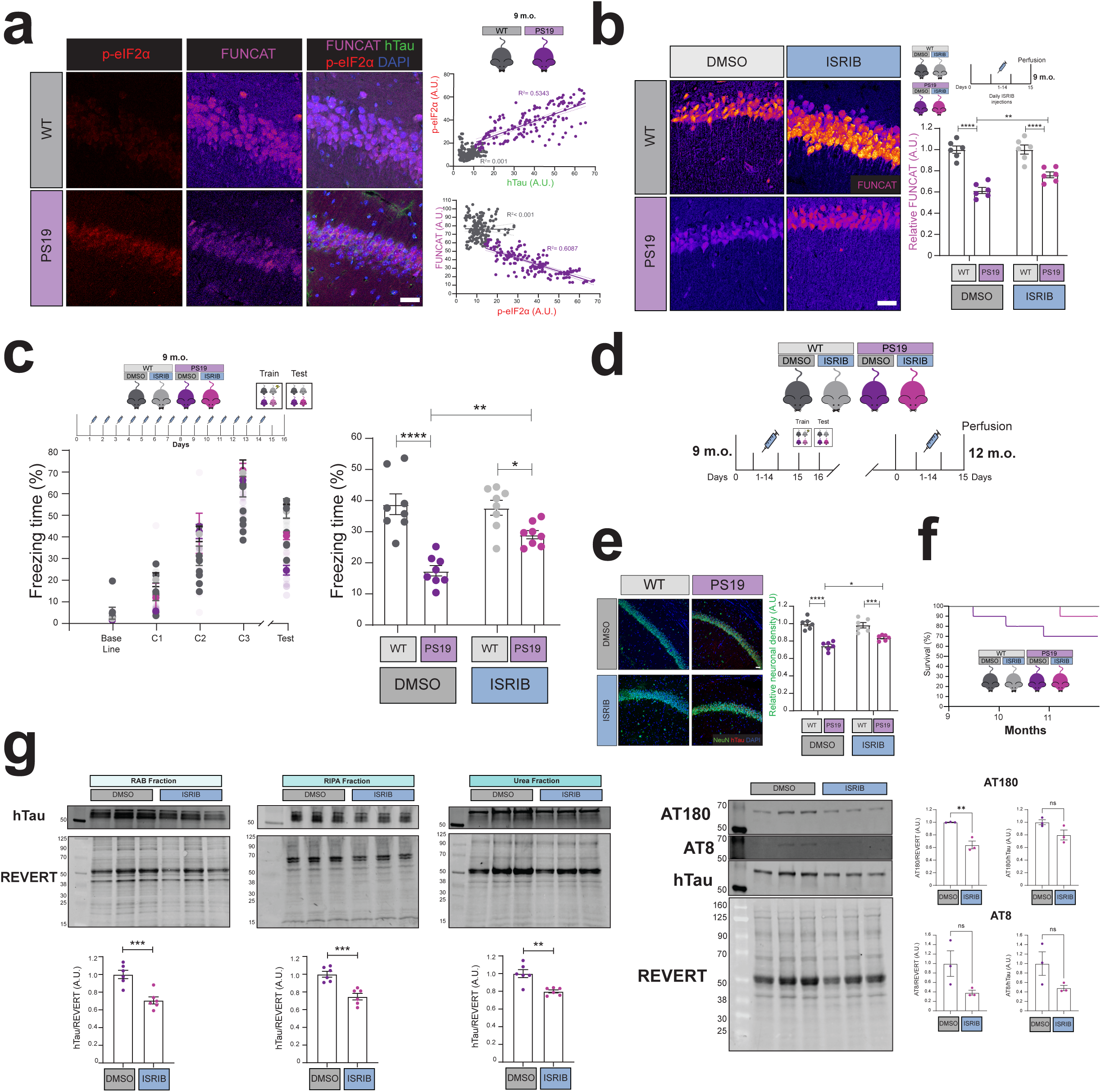
Preventing the integrated stress response rescues memory formation, attenuates neuronal death, decreases tau accumulation and phosphorylation, and improves survival in PS19 mice. (A) Increased p-eiF2α is correlated with increased human tau and decreased protein synthesis levels in the hippocampal neurons of 9-month-old PS19 mice (Pearson’s correlation, n=4 animals, data points = neurons). (B) Administration of the ISR inhibitor, ISRIB rescues homeostatic mRNA translation in the hippocampal neurons of 9-month-old PS19 mice (Two-way ANOVA, Tukey’s MCT, **=p≤0.01 ****=p≤0.0001, n=6 animals) Scale bar = 40 μm. (C) Pharmacological inhibition of the integrated stress response by ISRIB partially rescues long-term memory formation following contextual fear conditioning in 9 month-old PS19 mice (Two-way ANOVA, Tukey’s MCT, *=p≤0.05, **=p≤0.01, ****=p≤0.0001, n=8 animals, error bars=S.E.M.). (D) Schematic of repeated ISRIB treatment in PS19 mice. (E) Repeated administration of ISRIB partially attenuates hippocampal neuronal loss in 12-month-old PS19 mice (Two-way ANOVA, Tukey’s MCT, *=p≤0.05, **=p≤0.01, n≥6 animals, error bars=S.E.M.) Scale bar = 40 μm. (F) Preventing ISR activity through ISRIB improves the survival rates of PS19 mice at 12-months-old. (G) Repeated ISRIB treatment decreases the accumulation of total tau in the hippocampus of 12-month-old PS19 mice in RAB, RIPA, and Urea fractions (Welch’s T.test, n=6 animals, **=p≤0.01, ***=p≤0.001) as well as decreasing the total levels of AT8 & AT180 tau phosphorylation (Welch’s T.test, n=3 animals, *=p≤0.05, **=p≤0.01).

Next, we examined how repeated treatments with ISRIB altered neuronal loss in PS19 mice. After 9 months of age, PS19 mice start to experience neurodegeneration, characterized by a thinning of the CA1 layer of the hippocampus(Yoshiyama *et al*, 2007). We treated mice with ISRIB at both 9- and 12-months of age, before using NeuN staining to quantify the number of neurons in the CA1 layer of the hippocampus. Blocking ISR activity attenuated neuronal loss in PS19 mice, with ISRIB treated mice exhibiting a ≈15% loss in the number of CA1 hippocampal neurons compared to the ≈25% loss observed in the DMSO treated controls (Fig 4E). Repeated ISRIB treatment also appears to improve survival rates of PS19 mice, with ISRIB-treated mice experiencing a 10% mortality rate by 12 months of age, opposed to the 30% rate of DMSO-treated controls (Fig 4F).

Finally, we sought to determine whether repeated inhibition of the ISR decreased tau accumulation and phosphorylation. Biochemical analysis following sequential protein extraction found that PS19 mice treated with ISRIB exhibit reduced tau accumulation, regardless of solubility (Fig 4G). ISRIB treatment also reduced both AT180 and AT8 tau phosphorylation in 12-month-old PS19 mice in the hippocampus (Fig 4G).

## Discussion

Over the past decade a substantial body of work has suggested that the dysregulation of mRNA translation is a hallmark of neurodegenerative diseases(Storkebaum *et al*, 2023). Despite this progress, the precise mechanisms by which this dysregulation occur remain unclear, as does the role this dysregulation plays in disease progression. Here we detail a novel mechanism by which FTD-mutant tau impairs translation by slowing ribosomal elongation speed and inducing ribosomal collision early in disease progression. We show that this ribosomal collision activates the integrated stress response via GCN2, further impairing translation. Lastly, we reveal that this ISR-induced impairment in translation causes memory impairments in the PS19 mouse model of FTD, and exacerbates other forms of pathology, including tau accumulation and phosphorylation, neuronal loss, and increased mortality.

Numerous recent studies have highlighted the potential therapeutic benefits for targeting the integrated stress response (ISR) when treating neurodegenerative diseases. The ISR is thought to be activated in several neurodegenerative disorders such as AD with increased p-eIF2a and PKR activity being observed in post-mortem tissue of AD patients compared to age-matched controls (Chang *et al*, 2002). Studies in rodent models have also suggested that ISR activity is increased in other neurodegenerative diseases such as FTD (Luan *et al*, 2023), amyotrophic lateral sclerosis(Saxena *et al*, 2009), and Parkinson’s disease(Hoozemans *et al*, 2007). The role of ISR activity as a driver of pathology in models of these diseases has primarily been investigated at later disease stages, with inhibiting the ISR and its upstream kinases after the development of symptoms being shown to rescue behavioral and electrophysiological deficits. For example, blocking ISR activity via ISRIB has been demonstrated to rescue synapse function and memory in the APPswe/PS1ΔE9 model of AD and mice treated with toxic amyloid-β oligomers (Oliveira *et al*, 2021).

In the context of tauopathies, 6-month-old rTg4510 mice, which overexpress P301L FTD-mutant tau, exhibit increased phospho-PERK and p-eIF2α concurrently with impaired memory and hippocampal neuronal loss, with PERK inhibition ameliorating these phenotypes (Radford *et al*, 2015). Our findings are consistent with these previous studies, furthering the argument that the inhibition of mRNA translation via the ISR may drive pathology in these neurodegenerative diseases.

The mechanisms by which the ISR is activated are likely to differ across neurodegenerative diseases. Indeed, although we demonstrate that GCN2 activity causes ISR activation in 6-month-old PS19 mice, inhibiting GCN2 did not fully reduce eIF2α phosphorylation to WT levels, suggesting the other ISR pathways are likely active. One potential mechanism of this additional concurrent ISR activation in neurodegenerative diseases characterized by the presence of aggregation-prone proteins, including tauopathies, is the unfolded protein response (UPR). In the UPR, the accumulation of misfolded proteins can result in a chronic increase in ISR activity through the activation of PER(Costa-Mattioli & Walter, 2020). Our findings highlight the possibility that these different mechanisms of ISR activation are triggered at different stages of disease progression. Currently however, it is unknown whether the activation of the ISR via GCN2 observed in PS19 mice occurs independently of other forms of ISR activation or if these phenomena are interconnected. Furthermore, it is unclear if GCN2-ISR activation induced by ribosomal collisions is unique to tauopathies, or if this only recently described phenomenon occurs in other neurodegenerative diseases.

Interestingly, we found that FTD-mutant tau also inhibits mRNA translation via a mechanism completely independent of ISR activation. mRNA translation is significantly decreased in the PS19 mouse model of FTD months prior to the activation of the ISR, with ISR inhibition having no effect of mRNA translation levels at this early disease stage. Instead, we found that FTD-mutant tau slows ribosome elongation speed prior to the onset of the majority of FTD-like symptoms observed in these mice, such as impaired LTP, impaired memory, tangle formation, as well as synaptic and neuronal loss. One possible explanation for this is that FTD-mutant tau may directly disrupt mRNA translation, potentially through its interaction with ribosomes (Tracy *et al*, 2022; Banerjee *et al*, 2020). Indeed, some findings suggest that tau exhibits altered interactions with ribosomes in disease (Meier *et al*, 2016), and may disrupt the formation of polysomes (Evans *et al*, 2021b). However, although the FTD-mutant tau induced slowing of ribosomal elongation speed likely contributes to the ≈20% reduction in global translation observed in young PS19 mice, we cannot disregard other potential mechanisms by which translation may be impaired at this age, such as decreased mRNA and ribosome availability, altered metabolite levels, or aberrant signaling.

The mechanism by which FTD-mutant tau slows ribosomal elongation is currently unknown. It is also unclear if pathogenic tau slows ribosomal elongation speed for all mRNAs, or if only certain mRNAs are affected. One potential link between pathogenic tau and slowed ribosomal elongation is the mTORC1 pathway. The mTORC1 pathway regulates the expression of several components of machinery responsible for mRNA translation, including translation elongation factors (Hoeffer & Klann, 2010). Interestingly, the activity of the mTORC1 pathway is thought to be increased in tauopathies, with inhibition of mTORC1 by rapamycin being shown to attenuate tau accumulation, aggregation, and hyperphosphorylation in P301S-hTau expressing mice(Ozcelik *et al*, 2013; Suloh *et al*, 2025). Canonically, an increase in mTORC1 activity would be thought to boost ribosomal elongation speed by inhibiting eEF2K, and in turn, increasing the activity of eEF2. It is possible that the boosting of mTORC1 activity observed in tauopathies may serve as a means to compensate for the FTD-tau induced impairment in ribosomal elongation we describe here. Alternatively, pathogenic tau may interrupt mTORC1 signaling by modulating its downstream effectors, such as eEF2K, in turn slowing ribosomal elongation.

The slowing of ribosomal elongation speed observed in young PS19 mice may lead to ribosomal collision. Ribosomal collisions have been shown to occur in yeast and mammalian cells under conditions of UV stress(Sinha *et al*, 2024), amino acid deprivation(Zhou *et al*, 2025), and treatment with low doses of elongation inhibitors as well as alkylating and oxidizing agents (Wu *et al*, 2020). To the best of our knowledge, our study is the first to observe this phenomenon occurring in the mouse brain and the first to indicate that ribosomal collisions occur in the context of neurodegenerative disorders. Current techniques for the detection of ribosomal collision, including those used in this study, rely upon the inability of RNAses to digest the mRNA between two collided ribosomes. A caveat to this technique is the possibility that the disome peak observed is only due to incomplete digestion. Although this is unlikely given the relatively high concentration of RNAses used in our study, further characterization will be required to resolve the nature of these ribosomal collisions.

Interestingly, 3-month-old PS19 mice do not exhibit increased ISR activity despite presenting with increased ribosomal collisions compared to their WT counterparts. Although ribosomal collision can result in induction of the ISR via GCN2, this is only thought to occur if activation of the ribosomal quality control pathways is insufficient to resolve these collisions (Wu *et al*, 2020). It has been hypothesized ribosomal collision acts as “cellular gauge”, with cells modulating their response to ribotoxic insults depending on the prevalence of collided ribosomes. It is thought that low levels of collisions are resolved by the ribosomal quality control/no-go decay pathways, moderate levels of collisions activate GCN2 and the ISR, and high levels of unresolved collisions lead to apoptosis through activation of the stress-activated protein kinases p38 and JNK(Wu *et al*, 2020; Sinha *et al*, 2024). Given that ribosomal collision appears to be less pronounced in the 3-month-old PS19 mice compared to their 6-month-old counterparts, this “ribotoxic gauge” hypothesis may account for GCN2-mediated activation of the ISR only occurring after the continued accumulation of collided ribosomes in these older mice.

Given the reliance of neuronal plasticity and memory upon mRNA translation, it is surprising that the ≈20% reduction in homeostatic protein synthesis observed in our 3-month-old PS19 mice was not sufficient to prevent the formation of new long-term memories. Indeed, PS19 mice only exhibit impaired fear associative memory at later timepoints when mRNA translation is decreased by ≈40%, with our GCN2iB and ISRIB experiments demonstrating that these impairments in memory are caused, at least in part, by activation of the ISR. A simple explanation for this observation is that preventing ISR activity may preserve the neuron’s translational capacity to respond to stimuli. Both GCN2iB and ISRIB treatment restored homeostatic protein synthesis to ≈80% of WT levels, which is remarkably similar to the deficits in mRNA translation observed in young PS19 mice prior to ISR activation. It is possible that the combined effects of pathogenic tau both slowing ribosomal elongation speed, and subsequently impairing translation initiation, prevent neurons from synthesizing enough proteins to facilitate memory formation, and that relieving either of these proteostatic stressors would improve memory. A more nuanced alternative hypothesis to that of reduced translational capacity causing tau-induced memory impairment is that the activation of the ISR by pathogenic tau directly prevents memory-facilitating translation. A recent study highlighted the potential physiological role of the ISR in regulating memory-facilitating translation, with GADD34 rapidly dephosphorylating eIF2α following neuronal activity (Oliveira *et al*, 2024). It is possible that the GCN2-mediated activation of the ISR by pathogenic tau overwhelms the ability of GADD34 to dephosphorylate eIF2α during memory formation, in-turn preventing memory-associated and memory-facilitating translation.

Interestingly, despite returning homeostatic levels of protein synthesis to pre-ISR activation levels, neither GCN2iB or ISRIB treatment fully restored memory. Although it is not possible to discount the possibility that the remaining impairment in memory is due to pathogenic effects of tau independent of mRNA translation, one potential interacting factor worth exploring is aging. As cells age, they experience heightened stress, with a recent study demonstrating that the brains of aged killifish experience increased ribosomal stalling(Di Fraia *et al*, 2025). Therefore, although FTD-tau-induced slowing of ribosomal elongation was not sufficient to inhibit memory in 3-month-old PS19 mice, in older mice, the combination of this effect with the increased metabolic and proteostatic stress associated with aging may be sufficient to impair memory.

We also found that inhibition of the ISR attenuated both tau accumulation and phosphorylation. This is similar to studies in other neurodegenerative diseases, which found that inhibiting ISR activation can alter tau phosphorylation and may be neuroprotective (van der Harg *et al*, 2014; Lin *et al*, 2014; Radford *et al*, 2015). It is currently unknown how preventing ISR activity promotes the clearance of these aggregation-prone proteins, but one exciting potential explanation is a hypothesized link between protein synthesis and protein degradation. Indeed, a previous study has shown that inhibiting protein degradation leads to a decrease in protein synthesis, which is achieved through activation of the ISR by HRI (Alvarez-Castelao *et al*, 2020). We also found this to be the case in HEK293 cells, where we show that FTD-mutant tau decreases protein synthesis even when proteasome degradation is blocked through the addition of MG132 (Supplementary Fig 5). Although currently undescribed, it is possible that the inverse of this regulation occurs, with protein degradation levels being tied to protein synthesis levels in order to maintain proteostasis. Further research will be required to determine if this is the case, and if so, if this results in aggregation-bound proteins exhibiting slowed degradation when protein synthesis is impaired.

Numerous pharmaceutical trials are currently in preparation to assess the potential therapeutic value of molecules which target the ISR, such as ISRIB, in treating neurodegenerative diseases (Bravo-Jimenez *et al*, 2025). Our findings support this therapeutic approach, suggesting that ISR activation may cause the memory impairments observed in tauopathies such as FTD and exacerbate other aspects of tau pathology, such as tau accumulation, tau phosphorylation, and neuronal death. Despite this optimism, it is important to note that memory is both nuanced and complicated, and that the regulation of mRNA translation underpinning it is likely even more so. Although chronically decreasing the effects of p-eIF2α partially rescued memory formation in our mice, the accuracy of this memory is unclear. Studies which have artificially boosted protein synthesis in mice have shown that this can generalize memories (Shrestha *et al*, 2020a) and leads to improper memory extinction or behavioral inflexibility(Trinh *et al*, 2012). As such, it is possible that in order to be an effective treatment for neurodegenerative diseases in humans, treatments which seek to modulate mRNA translation regulatory pathways such as the ISR may need to do so in a more targeted and controllable way.

Collectively, our data adds to the growing body of evidence to suggest that the emerging hallmark of neurodegenerative diseases, dysregulated mRNA translation, is not just a consequence of disease, but may indeed drive it. We identify a novel mechanism where pathogenic tau slows translation elongation and causes ribosomal collision before impairing memory formation through GCN2-mediated activation of the ISR. Our findings detail how the mechanisms of mRNA translation dysregulation are altered throughout disease progression and demonstrate a mechanism by which impaired translation may play a causative role in driving memory impairments in neurodegenerative diseases.

## Methodology

### Animal Care

Male and female PS19 mice on the C57/BL6J background (Jackson Laboratories, 024841) and wild-type littermates were provided with both food and water *ad libitum* and maintained on a 12h/12h light/dark cycle at a stable temperature (78°F) and humidity (40–50%). All procedures involving the use of animals were performed in accordance with the guidelines of the National Institutes of Health and were approved by the Institutional Animal Care and Use Committee (IACUC, protocols 221-1143 and 221-1145). A roughly even distribution of male and female mice was used across groups in all experiments.

### Labelling of newly synthesized proteins with azidohomoalanine in awake mice

Azidohomoalanine (Vector Laboratories, CCT-1106) was delivered to awake mice via retro-orbital injection as previously described (Evans *et al*, 2025). In brief, a 0.5% proparacaine hydrochloride ophthalmic solution (Covetrus, 2963726) was first administered to the eye 2 minutes prior to RO injection. 50 mg/kg AHA dissolved in phosphate-buffered saline (PBS) was then injected retro-orbitally by two researchers with the first restraining the mouse’s head before subsequently drawing back the skin below the eye and the second researcher using a 27G needle to inject into the retro-bulbar sinus.

Two hours following AHA injection, mice were deeply anesthetized with isoflurane before being intracardially perfused with 20 mL of PBS. For FUNCAT-WB analysis, the hippocampus was then dissected before being snap-frozen. For FUNCAT/IHC analysis, one hemisphere was fixed with 4% paraformaldehyde (PFA, ThermoFisher, 50-980-494).

### Contextual fear conditioning

All rodent behavioral training and analysis was performed during the light cycle of the mice, with mice being handled daily for at least 7 days prior to training. Contextual fear conditioning was performed as previously described (Evans *et al*, 2025). In brief, mice were placed into a fear conditioning chamber (Coulbourn instruments) for 270 seconds before receiving a 2 second, 0.5mA foot-shock, with this foot-shock being repeated 100 and 200 seconds later. The mouse was then removed from the chamber 150 seconds after the administration of the final shock, spending a total of 12 minutes in the chamber. No shock control mice were placed in the chamber for the same period of time but did not receive a foot shock.

To label memory-associated protein synthesis in these mice, AHA was delivered via RO injection immediately following training, with mice being perfused two hours later.

To test the ability of mice to recall the association between the context and receiving the foot shock, a probe test was conducted 24 hours following training, where mice were placed in the context for 300 seconds without receiving a foot shock. Freezing behavior was automatically measured by Freeze Frame 4 software (ActiMetrics).

### ISRIB and GCN2iB treatment in mice

To inhibit GCN2 activity in mice, PS19 mice and age-matched WT controls were injected intraperitoneally daily for 14 days with 5 mg/kg of GCN2iB (MedChemExpress, HY-112654) dissolved in DMSO and diluted in PBS. Contextual fear conditioning was conducted one day after the final GCN2iB injection, followed by AHA treatment one day after the probe trial.

To determine if ISRIB treatment rescued mRNA translation in PS19 mice, mice were intraperitoneally injected daily for 14 days with 0.25 mg/kg of ISRIB (Sigma Aldrich, SML0843) dissolved in DMSO and diluted in PBS as previously described (Oliveira *et al*, 2021). Mice were retro-orbitally injected with AHA one day after the final ISRIB treatment. To determine if ISRIB treatment rescued memory in PS19 mice, mice were given the same treatment paradigm as above, with contextual fear conditioning being conducted one day after the final ISRIB injection. These mice were then tracked between the ages of 9 and 12 months to examine the effects of ISRIB treatment on survival, and then once again treated with ISRIB to examine the effects of repeated ISRIB treatment on tau pathology.

### Immunohistochemistry and FUNCAT staining

To visualize AHA labeled proteins in the brains of PS19 and WT mice, fixed brains were submerged in PBS with 30% w/v sucrose for 48 hours prior to being sectioned at 40 μM thickness using a vibratome (Leica, VT1200). Samples were placed in 5% bovine serum albumin, 5% normal goat serum, and 0.5% Triton-x in PBS for 1h at RT under constant agitation. 5μM Alexa 647 alkyne (Vector Laboratories, CCT-1301) was then used in combination with the Click-&-Go® Cell Reaction Buffer Kit (Vector Laboratories, CCT-1263) to fluorescently label newly synthesized proteins as per the manufacturer’s instructions. Neurons were visualized using a guinea pig IgG anti-NeuN primary antibody (Synaptic Systems, 266 004, 1:1000). hTau was visualized using the mouse primary antibody Tau12 (Sigma Aldrich, MAB2241, 1:2000) and p-eIF2α was visualized using a rabbit anti-p-eIF2α (Ser51) primary antibody (Cell Signaling, 9721, 1:500). Primary antibodies were visualized using Alexa-405 labeled goat anti-Guinea Pig IgG (H+L) (Abcam, ab175678), Alexa-488 labeled goat anti-Mouse IgG (H+L) (Invitrogen, A-11001), and Alexa-594 labeled goat anti-Rabbit IgG (H+L) (Invitrogen, A-11012) secondary antibodies. DAPI was used to stain cell nuclei and samples were mounted in ProLong Gold mounting media (ThermoFisher, P10144).

### Cell Culture

HEK293 cells were cultured in Dulbecco’s modified Eagle’s medium (DMEM) (Gibco, 21013024) supplemented with 10% fetal bovine serum and 50 U/ml penicillin/streptomycin at 37 °C in a 5% CO_2_ saturated humidity incubator. Cells were then transfected with either non-mutant, P301S, or V337M mutant full length human tau or an emerald fluorescent protein control using lipofectamine LTX (Thermo Fisher, 15,338,100), as per the manufacturer’s instructions. These constructs were FLAG-tagged and in the pCMV backbone (Addgene, 11153).

For the labelling of newly synthesized proteins, cells were treated with 1mM AHA for 4 hours. In experiments where proteasome activity was blocked, cells were treated with 10μM MG-132 (Sigma Aldrich, M8699).

### Analysis of ribosomal elongation speed via harringtonine run-off assays

Harringtonine run-off assays in HEK293 cells were performed as previously described (Mohamed & Klann, 2023), with minor modifications. Briefly, cells were treated with 2μg/mL harringtonine (Abcam, ab141941) prior to being treated with 5μg/mL puromycin (Gibco, A1113803) for 15 minutes. As a negative control, cells were treated with 200 mM emetine (Millipore, 324693), an inhibitor of translation elongation. Puromycin incorporation into nascent poly-peptide chains was then detected using a mouse anti-puromycin primary antibody (Sigma-Aldrich, MABE343, 1:1000) and Alexa-488 labeled goat anti-Mouse IgG (H+L) (Invitrogen, A-11001) secondary antibody. Fluorescence measurements were made on a SpectraMax ID3 (Molecular Devices) with the total puromycin signal being normalized to DAPI fluorescence.

For harringtonine run-off assays in mice, live hippocampal slices from 3-month-old PS19 mice and wild-type littermates were prepared as previously described (Evans *et al*, 2024). After recovery, these slices were treated with 1 mM AHA for 30 minutes before being treated with 5μg/mL harringtonine for varying amounts of time. All sections were treated with AHA for a total of 1 hour.

### Biochemical and FUNCAT-Western Blot analysis

Proteins were extracted from snap-frozen brain samples, hippocampal slices, and resuspended cells, via sonication. For sequential extraction, samples were first placed in reassembly (RAB) buffer (G-Biosciences, 786-91) with Halt protein inhibitor cocktail (ThermoFisher, 78,438) and 1 mM phenylmethylsulfonyl fluoride (PMSF) (ThermoFisher, 36,978). Samples were then centrifuged at 21,000xg for 90 minutes at 4°C, with supernatant being collected. The remaining pellet was then sonicated in radioimmunoprecipitation assay (RIPA) buffer (Cell Signaling, 9806) with Halt and 1 mM PMSF, before the centrifugation was repeated and supernatant collected. Lastly, this remaining pellet was sonicated in 8M Urea (Millipore Sigma, U4883) in 500mM Tris-HCl pH 8.5 with Halt and 1mM PMSF before centrifugation and supernatant collection. For all other experiments, proteins were extracted in RIPA buffer with Halt and 1mM PMSF. The EZQ protein quantification assay (Invitrogen, R33200) was used to determine protein concentration following manufacturers instructions.

Samples were denatured by boiling at 95°C for 1 minute in 1X Bolt™ LDS Sample Buffer (Invitrogen, B0007) and 0.1 M DTT (Millipore Sigma, 646563). Samples were then separated via SDS-PAGE using 4-12% Bis-Tis gels (Invitrogen, NW04127BOX) before being transferred to a PVDF membrane (Invitrogen, IB24002) using the iBlot semidry transfer system (Invitrogen, IB2100). For the detection of GCN2 and p-GCN2, 3-10% Tris-acetate gels (Invitrogen, EA0378BOX) were used. The total protein stain REVERT (LI-COR, 926–10,011) was used for normalization.

p-GCN2 was detected using a rabbit anti-p-GCN2 (T889) primary antibody (Abcam, ab75836, 1:500). GCN2 was detected using a mouse anti-total-GCN2 primary antibody (SantaCruz Biotechnology, 1:500). eIF4B was detected using a rabbit anti-eIF4B primary antibody (Cell Signaling, 3592, 1:1000). p-eIF2a was detected using a rabbit anti-p-eIF2α (Ser51) primary antibody (Cell Signaling, D9G8, 1:1000). Total eiF2a was detected using a mouse anti-eIF2α primary antibody (Cell Signaling, 2103, 1:1000). These primary antibodies were visualized using HRP-conjugated Anti-Rabbit IgG (H+L) (Promega, W4011) or HRP-conjugated Anti-Mouse IgG (H+L) (Promega, W4021). Human tau was detected using Tau 12 (1:2000) and IRDye® 680RD Goat Anti-Mouse IgG (LI-COR, 926-68070, 1:10000). Restore™ Western Blot Stripping Buffer (ThermoFisher, 21059) was used to strip HRP staining between primary antibody stains.

For the detection of AHA-labelled proteins, samples were labelled with IRDye® 800CW Alkyne Infrared Dye (LI-COR, 929-60002) using the Click-&-Go® Protein Reaction Buffer Kit (Vector Laboratories, CCT-1262) as previously described (Evans *et al*, 2025). These samples were then ran on separate blots as FUNCAT labelling can interfere with antibody binding.

### Polysome profiling and detection of collided ribosomes

To extract samples for polysome profiling, mice were first deeply anesthetized with isoflurane before being euthanized via decapitation with the hippocampus then being dissected and snap frozen in liquid nitrogen. Working in RNAse-free conditions, samples were then lysed on ice in polysome lysis buffer (20mM Tris pH 7.5, 150 mM NaCL, 5mM MgCl_2_, 1mM DTT, 200 μg/mL cycloheximide (Sigma Aldrich, 01810), 1X Halt, 24U/mL RQ1 DNAse (Promega, M6101) and 1% triton-x in H_2_O) via sonication. For HEK293 cells, cells were first incubated with 200 μg/mL cycloheximide for 3 minutes a 37°C, before being washed with ice-cold PBS with 100 μg/mL cycloheximide and then lysed in polysome lysis buffer.

All samples were centrifuged for 10 minutes at 13,000xg at 4°C. RNA concentration was quantified via the RediPlate 96 RiboGreen RNA quantification assay kit (Invitrogen, R32700) as per the manufacturer’s instructions. Samples were then digested with 37.5 U/1000ng RNA of RNAse A (Invitrogen, AM2271) and 9.375 U/1000ng RNA of RNAse T1 (ThermoFisher, EN0542) for 20 minutes at RT. RNAse activity was then quenched through the addition of 400U/mL of RNasin™ Plus RNase Inhibitor (Promega, N2615). Samples were then separated on a 10-50% linear sucrose gradient by centrifugation at 36,000xg using a SW41Ti rotor for 2 hours and 45 minutes at 4°C. The absorbance at 260nm across this gradient was then measured using a BioCOMP TRIAX gradient profiler, with the area under the curve of the monosome peak being used for normalization. Fractions were collected and proteins were extracted using chloroform-methanol precipitation. Extracted proteins were probed using the western blot protocol detailed above.

### Imaging and image analysis

For western blot analysis, fluorescently labelled membranes were imaged using a LI-COR Odessey M Scanner with the LI-COR Emperia Studio software being used for quantification. HRP-stained membranes were imaged using the FluorChem E System (ProteinSimple) and quantified using AlphaView.

For FUNCAT-IHC analysis, 15 μm thick Z-stack images were taken using a Leica SP8 Confocal microscope with maximum intensity projections being created in ImageJ. Image analysis was performed blinded in ImageJ with NeuN immunoreactivity being used to generate a mask. Mean gray value was then measured within this neuronal mask for each image, with no significant difference being detected between the area of these masks across groups. For the per neuron analysis of FUNCAT, hTau and p-eIF2α levels, ROIs were drawn around individual neurons using the NeuN stain. For the quantification of hippocampal neuron density, neurons were counted using the analyze particles plugin of ImageJ and normalized to the area of the granular layer of the CA1.

### Statistical analysis

GraphPad Prism 10.1.2 was used for statistical analysis, with a two-way ANOVA with Tukey’s multiple comparison test (MCT), Sidak’s MCT, Welch’s T.Test, Spearman’s correlation, or Pearson’s correlation being used as appropriate. All values are given as mean ± standard error of the mean. Significance was defined as *p < 0.05, **p < 0.01, ***p < 0.001, ****p < 0.0001.

**Supplementary Figure 1:**
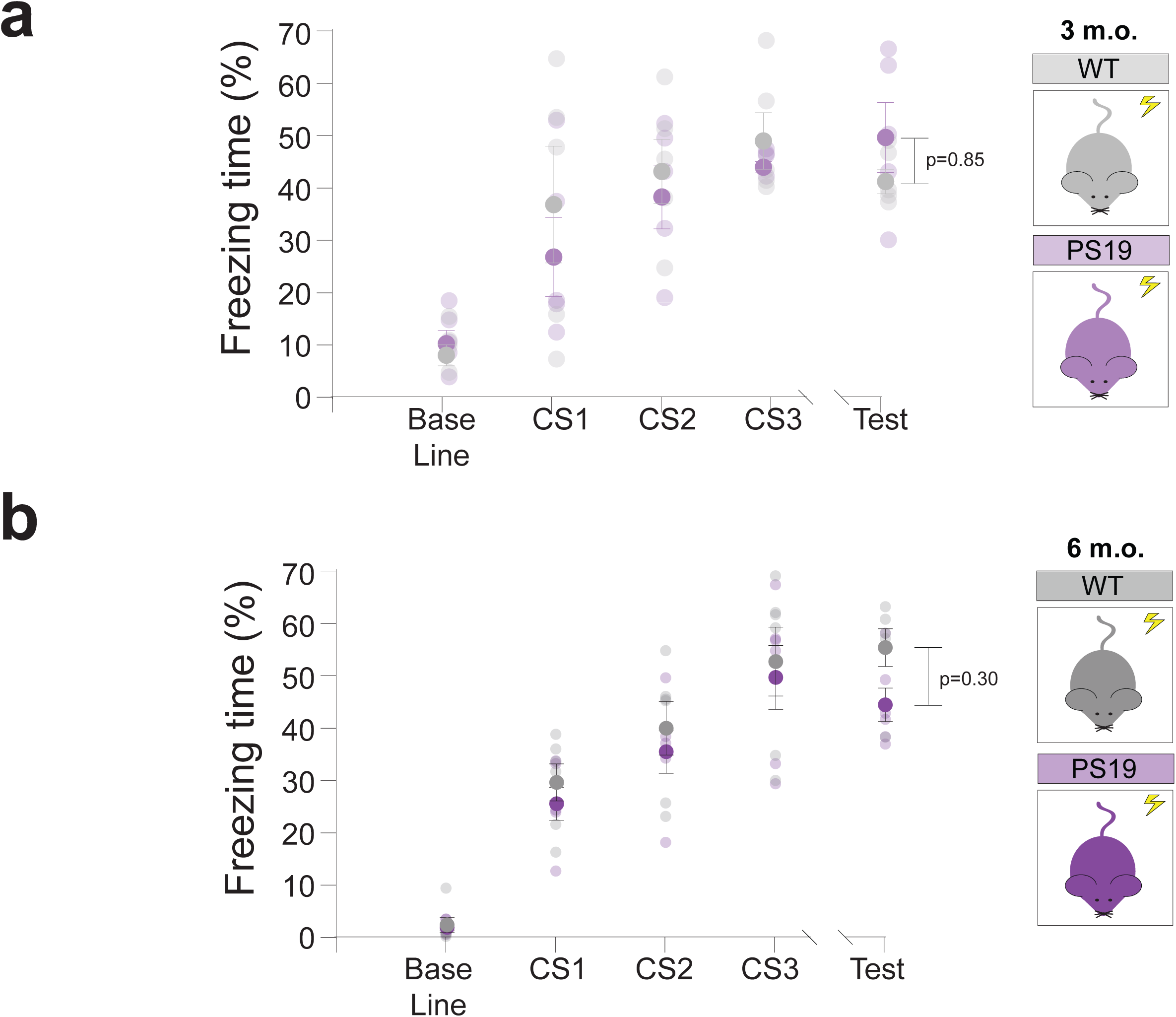
Contextual fear conditioning is not impaired in young PS19 mice. (A) 3-month-old PS19 mice freeze at similar rates to WT controls in a probe test 24 hours following training (Two-way ANOVA, Sidak’s MCT n= 5 animals). (B) 6-month-old PS19 mice exhibit a trend towards decreased freezing compared to WTs in the probe trial but this does not reach significance (Two-way ANOVA, Sidak’s MCT n= 6 animals).

**Supplementary Figure 2:**
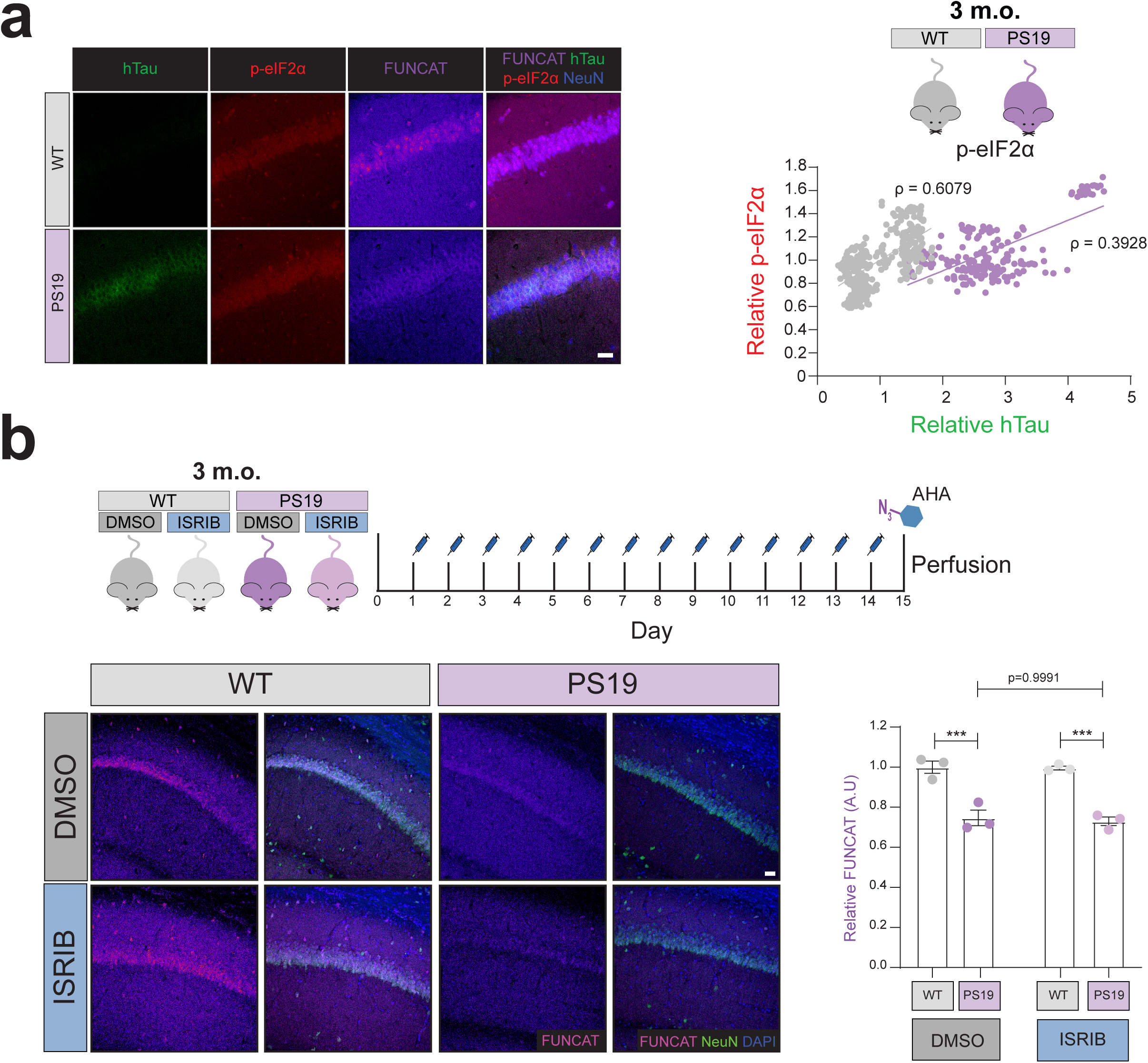
The integrated stress response is not activated in 3-month-old PS19 mice. (A) Total levels of p-eIF2a are not increased in the hippocampal neurons of 3-month-old PS19 mice compared to WT controls (Spearman’s correlation, n=6 animals, data points = neurons). (B) Daily i.p. administration of the ISR inhibitor, ISRIB, for 14 days, does not rescue homeostatic protein synthesis levels in 3-month-old PS19 mice (Two-way ANOVA, Tukey’s MCT, n=3 animals,***=p≤ 0.001). Scale bar = 40 μm.

**Supplementary Figure 3:**
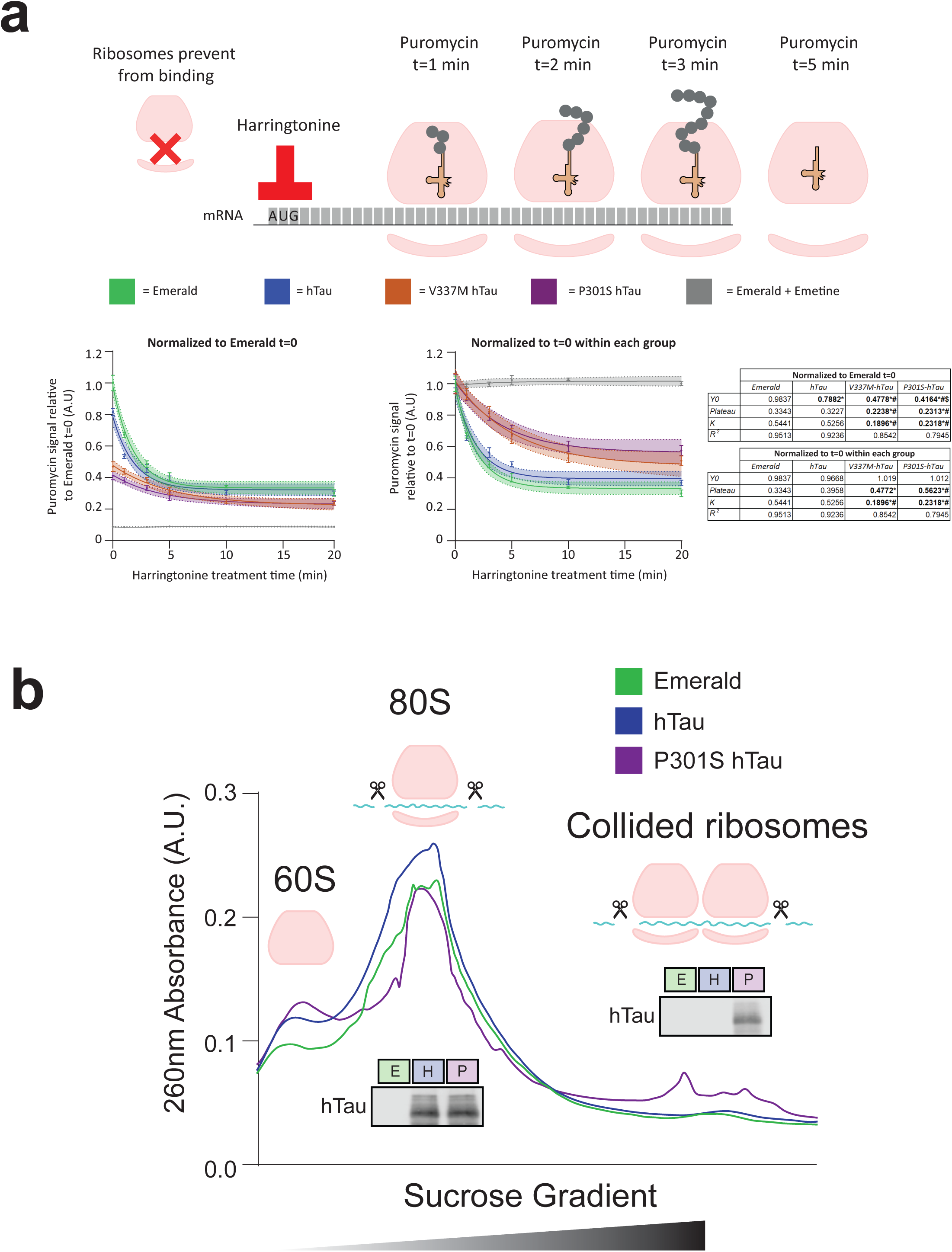
FTD-mutant tau expression in HEK293 cells recapitulates the results observed in PS19 mice. (A) Harringtonine run-off assays reveal that both P301S and V337M full length human tau slows ribosomal elongation speed compared to non-mutant full length human tau and a fluorescent protein control (Single-phase logarithmic decay, ∑ of squares F-test, 95% confidence interval is shown shaded, n=4, *= p<0.05 to Emerald, # = p<0.05 to hTau). (B) Averaged polysome profiles of RNAse digested HEK293 cell lysate demonstrates that P301S-hTau expression increased ribosomal collision compared to non-mutant hTau and emerald, a fluorescent protein control (n=4).

**Supplementary Figure 4:**
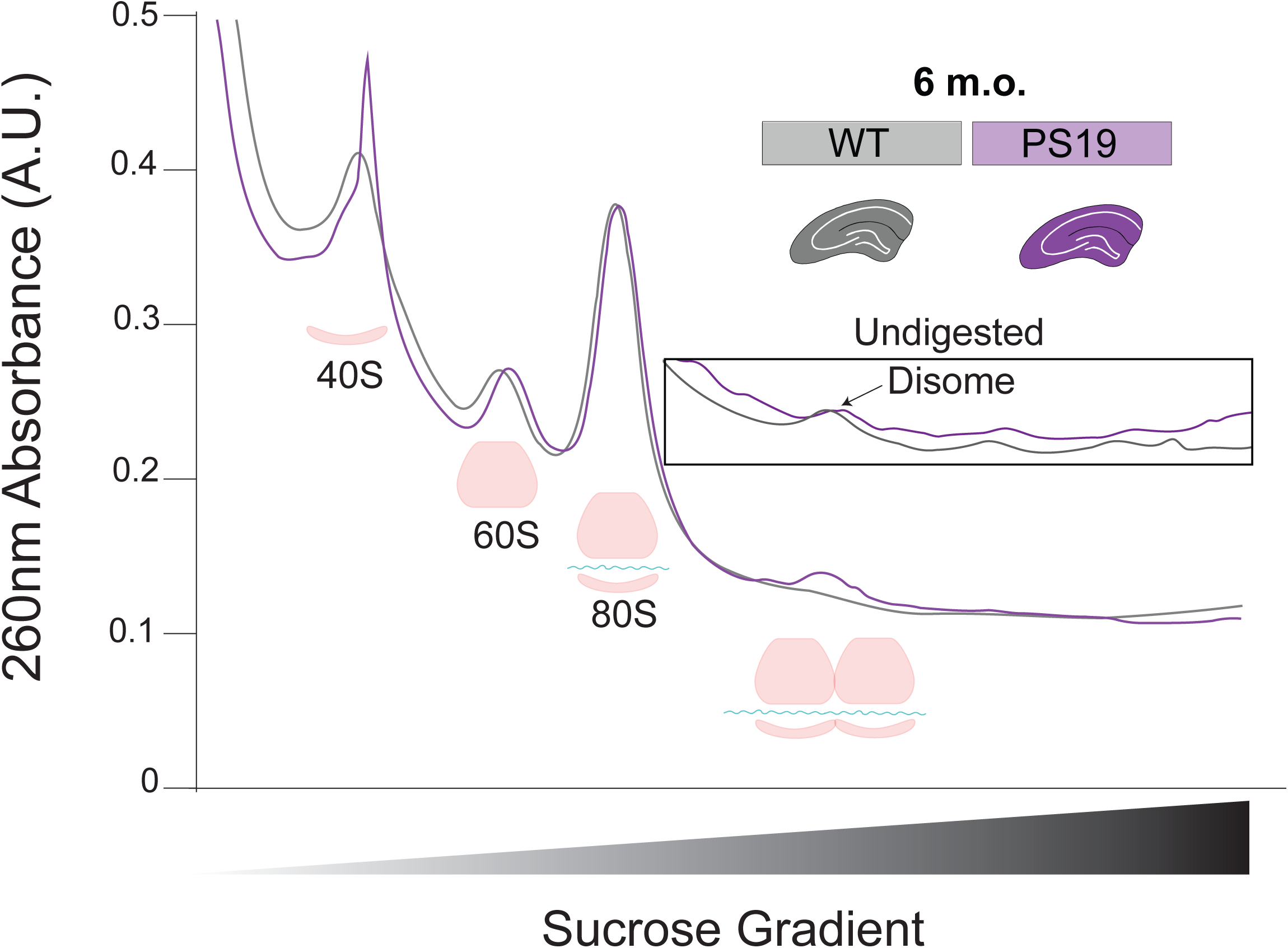
Polysome profiling following RNAse digestion reveals ribosomal collision in PS19 mice. Averaged polysome profiles of RNAse digested hippocampal lysate taken from 6-month-old PS19 mice (n≥5 mice).

**Supplementary Figure 5:**
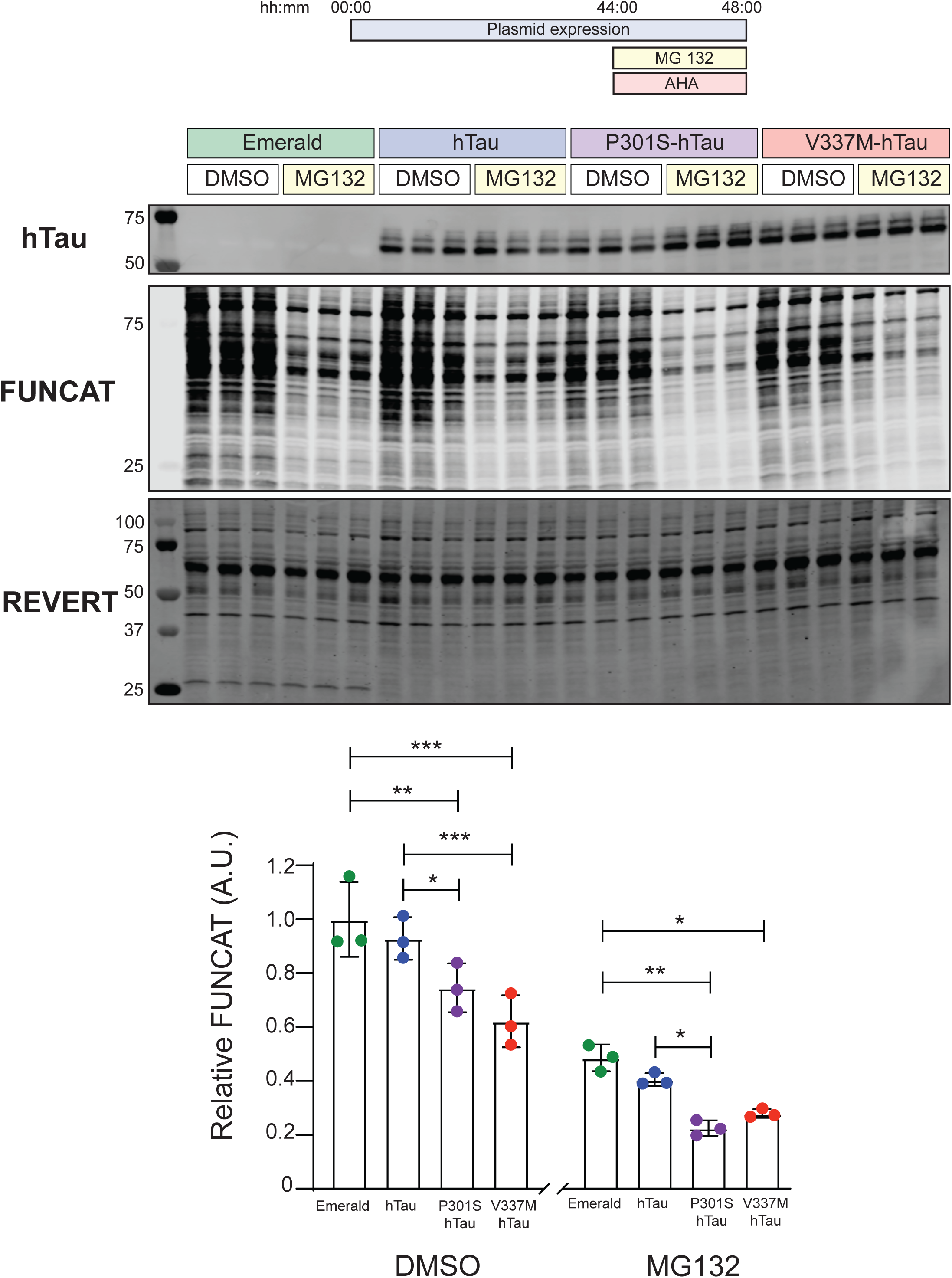
FTD-mutant hTau decreases protein synthesis in HEK293 cells, even in the presence of the proteasome inhibitor MG132

## Acknowledgement

The authors would like to acknowledge the contributions of Maggie Donohue and Jenesha Rawlani to this work.

## References

1. Alvarez-Castelao B, tom Dieck S, Fusco CM, Donlin-Asp P, Perez JD & Schuman EM (2020) The switch-like expression of heme-regulated kinase 1 mediates neuronal proteostasis following proteasome inhibition. Elife 9: 1–26

2. Apicco DJ, Ash PEA, Maziuk B, LeBlang C, Medalla M, Al Abdullatif A, Ferragud A, Botelho E, Ballance HI, Dhawan U, et al (2018) Reducing the RNA binding protein TIA1 protects against tau-mediated neurodegeneration in vivo. Nat Neurosci 21: 72–80

3. Banerjee S, Ferdosh S, Ghosh AN & Barat C (2020) Tau protein-induced sequestration of the eukaryotic ribosome: Implications in neurodegenerative disease. Sci Rep 10: 5225

4. Biever A, Glock C, Tushev G, Ciirdaeva E, Dalmay T, Langer JD & Schuman EM (2020) Monosomes actively translate synaptic mRNAs in neuronal processes. Science (1979) 367

5. Bowles KR, Silva MC, Whitney K, Bertucci T, Berlind JE, Lai JD, Garza JC, Boles NC, Mahali S, Strang KH, et al (2021) ELAVL4, splicing, and glutamatergic dysfunction precede neuron loss in MAPT mutation cerebral organoids. Cell 184: 4547–4563.e17

6. Bravo-Jimenez MA, Sharma S & Karimi-Abdolrezaee S (2025) The integrated stress response in neurodegenerative diseases. Mol Neurodegener 20: 20

7. Caccamo A, Majumder S, Richardson A, Strong R & Oddo S (2010) Molecular Interplay between Mammalian Target of Rapamycin (mTOR), Amyloid-β, and Tau. Journal of Biological Chemistry 285: 13107–13120

8. Chang RCC, Wong AKY, Ng H-K & Hugon J (2002) Phosphorylation of eukaryotic initiation factor-2alpha (eIF2alpha) is associated with neuronal degeneration in Alzheimer’s disease. Neuroreport 13: 2429–32

9. Costa-Mattioli M & Walter P (2020) The integrated stress response: From mechanism to disease. Science (1979) 368

10. Creekmore BC, Watanabe R & Lee EB (2024) Neurodegenerative Disease Tauopathies. Annual Review of Pathology: Mechanisms of Disease 19: 345–370

11. Davis HP & Squire LR (1984) Protein synthesis and memory: A review. Psychol Bull 96: 518–559

12. Ding Q, Markesbery WR, Chen Q, Li F & Keller JN (2005) Ribosome Dysfunction Is an Early Event in Alzheimer’s Disease. The Journal of Neuroscience 25: 9171–9175

13. Dubue JD, McKinney TL, Treit D & Dickson CT (2015) Intrahippocampal Anisomycin Impairs Spatial Performance on the Morris Water Maze. Journal of Neuroscience 35: 11118–11124

14. Elder MK, Erdjument-Bromage H, Oliveira MM, Mamcarz M, Neubert TA & Klann E (2021) Age-dependent shift in the de novo proteome accompanies pathogenesis in an Alzheimer’s disease mouse model. Commun Biol 4: 823

15. Evans HT, Adler D, Selles MC, Kalavai S V., Yu A, Wu V, Denil E, Golhan EN, Polavarapu A, Balamoti E, et al (2025) Mapping the spatiotemporal dynamics of de novo protein synthesis during long-term memory formation. doi:10.1101/2025.04.17.649250 [PREPRINT]

16. Evans HT, Benetatos J, van Roijen M, Bodea L-G & Götz J (2019) Decreased synthesis of ribosomal proteins in tauopathy revealed by non-canonical amino acid labelling. EMBO J 38: e101174

17. Evans HT, Blackmore D, Götz J & Bodea LG (2021a) De novo proteomic methods for examining the molecular mechanisms underpinning long-term memory. Brain Res Bull 169: 94–103

18. Evans HT, Ko T, Oliveira MM, Yu A, Kalavai S V., Golhan EN, Polavarapu A, Balamoti E, Wu V, Klann E, et al (2024) Light-Activatable, Cell-Type Specific Labeling of the Nascent Proteome. ACS Chem Neurosci 15: 3473–3481

19. Evans HT, Taylor D, Kneynsberg A, Bodea L-G & Götz J (2021b) Altered ribosomal function and protein synthesis caused by tau. Acta Neuropathol Commun 9: 110

20. Fortin DA, Srivastava T, Dwarakanath D, Pierre P, Nygaard S, Derkach VA & Soderling TR (2012) Brain-Derived Neurotrophic Factor Activation of CaM-Kinase Kinase via Transient Receptor Potential Canonical Channels Induces the Translation and Synaptic Incorporation of GluA1-Containing Calcium-Permeable AMPA Receptors. Journal of Neuroscience 32: 8127–8137

21. Di Fraia D, Marino A, Lee JH, Kelmer Sacramento E, Baumgart M, Bagnoli S, Balla T, Schalk F, Kamrad S, Guan R, et al (2025) Altered translation elongation contributes to key hallmarks of aging in the killifish brain. Science (1979) 389

22. Fresno M, Jiménez A & Vázquez D (1977) Inhibition of translation in eukaryotic systems by harringtonine. Eur J Biochem 72: 323–30

23. Fusco CM, Desch K, Dörrbaum AR, Wang M, Staab A, Chan ICW, Vail E, Villeri V, Langer JD & Schuman EM (2021) Neuronal ribosomes exhibit dynamic and context-dependent exchange of ribosomal proteins. Nat Commun 12: 6127

24. Gistelinck M, Lambert J-C, Callaerts P, Dermaut B & Dourlen P (2012) Drosophila Models of Tauopathies: What Have We Learned? Int J Alzheimers Dis 2012: 1–14

25. Götz J, Bodea L-G & Goedert M (2018) Rodent models for Alzheimer disease. Nat Rev Neurosci 19: 583–598

26. Götz J, Halliday G & Nisbet RM (2019) Molecular Pathogenesis of the Tauopathies. Annual Review of Pathology: Mechanisms of Disease 14: 239–261

27. van der Harg JM, Nölle A, Zwart R, Boerema AS, van Haastert ES, Strijkstra AM, Hoozemans JJ & Scheper W (2014) The unfolded protein response mediates reversible tau phosphorylation induced by metabolic stress. Cell Death Dis 5: e1393–e1393

28. Hoeffer CA & Klann E (2010) mTOR signaling: At the crossroads of plasticity, memory and disease. Trends Neurosci 33: 67–75

29. Hoozemans JJM, van Haastert ES, Eikelenboom P, de Vos RAI, Rozemuller JM & Scheper W (2007) Activation of the unfolded protein response in Parkinson’s disease. Biochem Biophys Res Commun 354: 707–711

30. Kapur M, Monaghan CE & Ackerman SL (2017) Regulation of mRNA Translation in Neurons—A Matter of Life and Death. Neuron 96: 616–637

31. Kauwe G, Lokitiyakul D, Wong IL, Schneider K, Sridhar V, Sellegounder D, Ngwala YY, Yao L, Chen JH, Pareja-Navarro KA, et al (2025) Pathogenic tau inhibits synaptic plasticity by blocking eIF4B-mediated local protein synthesis. doi:10.1101/2025.09.11.675671 [PREPRINT]

32. Khan MR, Yin X, Kang S-U, Mitra J, Wang H, Ryu T, Brahmachari S, Karuppagounder SS, Kimura Y, Jhaldiyal A, et al (2023) Enhanced mTORC1 signaling and protein synthesis in pathologic α-synuclein cellular and animal models of Parkinson’s disease. Sci Transl Med 15

33. Koren SA, Galvis-Escobar S & Abisambra JF (2020) Tau-mediated dysregulation of RNA: Evidence for a common molecular mechanism of toxicity in frontotemporal dementia and other tauopathies. Neurobiol Dis 141: 104939

34. Koren SA, Hamm MJ, Meier SE, Weiss BE, Nation GK, Chishti EA, Arango JP, Chen J, Zhu H, Blalock EM, et al (2019) Tau drives translational selectivity by interacting with ribosomal proteins. Acta Neuropathol 137: 571–583

35. Lasagna-Reeves CA, de Haro M, Hao S, Park J, Rousseaux MWC, Al-Ramahi I, Jafar-Nejad P, Vilanova-Velez L, See L, De Maio A, et al (2016) Reduction of Nuak1 Decreases Tau and Reverses Phenotypes in a Tauopathy Mouse Model. Neuron 92: 407–418

36. Lin L, Yang S-S, Chu J, Wang L, Ning L-N, Zhang T, Jiang Q, Tian Q & Wang J-Z (2014) Region-Specific Expression of Tau, Amyloid-β Protein Precursor, and Synaptic Proteins at Physiological Condition or Under Endoplasmic Reticulum Stress in Rats. Journal of Alzheimer’s Disease 41: 1149–1163

37. Lourenco MV, Clarke JR, Frozza RL, Bomfim TR, Forny-Germano L, Batista AF, Sathler LB, Brito-Moreira J, Amaral OB, Silva CA, et al (2013) TNF-α Mediates PKR-Dependent Memory Impairment and Brain IRS-1 Inhibition Induced by Alzheimer’s β-Amyloid Oligomers in Mice and Monkeys. Cell Metab 18: 831–843

38. Luan W, Wright AL, Brown-Wright H, Le S, San Gil R, Madrid San Martin L, Ling K, Jafar-Nejad P, Rigo F & Walker AK (2023) Early activation of cellular stress and death pathways caused by cytoplasmic TDP-43 in the rNLS8 mouse model of ALS and FTD. Mol Psychiatry 28: 2445–2461

39. Meier S, Bell M, Lyons DN, Rodriguez-Rivera J, Ingram A, Fontaine SN, Mechas E, Chen J, Wolozin B, LeVine H, et al (2016) Pathological Tau Promotes Neuronal Damage by Impairing Ribosomal Function and Decreasing Protein Synthesis. The Journal of Neuroscience 36: 1001–1007

40. Mohamed MS & Klann E (2023) Autism- and epilepsy-associated EEF1A2 mutations lead to translational dysfunction and altered actin bundling. Proceedings of the National Academy of Sciences 120

41. Moy JK, Khoutorsky A, Asiedu MN, Dussor G & Price TJ (2018) eIF4E Phosphorylation Influences Bdnf mRNA Translation in Mouse Dorsal Root Ganglion Neurons. Front Cell Neurosci 12

42. Nakamura A, Nambu T, Ebara S, Hasegawa Y, Toyoshima K, Tsuchiya Y, Tomita D, Fujimoto J, Kurasawa O, Takahara C, et al (2018) Inhibition of GCN2 sensitizes ASNS-low cancer cells to asparaginase by disrupting the amino acid response. Proceedings of the National Academy of Sciences 115: E7776–E7785

43. Natale C, Barzago MM & Diomede L (2020) Caenorhabditis elegans Models to Investigate the Mechanisms Underlying Tau Toxicity in Tauopathies. Brain Sci 10: 838

44. Oliveira MM, Lourenco M V., Longo F, Kasica NP, Yang W, Ureta G, Ferreira DDP, Mendonça PHJ, Bernales S, Ma T, et al (2021) Correction of eIF2-dependent defects in brain protein synthesis, synaptic plasticity, and memory in mouse models of Alzheimer’s disease. Sci Signal 14: eabc5429

45. Oliveira MM, Mohamed M, Elder MK, Banegas-Morales K, Mamcarz M, Lu EH, Golhan EAN, Navrange N, Chatterjee S, Abel T, et al (2024) The integrated stress response effector GADD34 is repurposed by neurons to promote stimulus-induced translation. Cell Rep 43: 113670

46. Ozcelik S, Fraser G, Castets P, Schaeffer V, Skachokova Z, Breu K, Clavaguera F, Sinnreich M, Kappos L, Goedert M, et al (2013) Rapamycin Attenuates the Progression of Tau Pathology in P301S Tau Transgenic Mice. PLoS One 8

47. Pisciottani A, Croci L, Lauria F, Marullo C, Savino E, Ambrosi A, Podini P, Marchioretto M, Casoni F, Cremona O, et al (2023) Neuronal models of TDP-43 proteinopathy display reduced axonal translation, increased oxidative stress, and defective exocytosis. Front Cell Neurosci 17

48. Radford H, Moreno JA, Verity N, Halliday M & Mallucci GR (2015) PERK inhibition prevents tau-mediated neurodegeneration in a mouse model of frontotemporal dementia. Acta Neuropathol 130: 633–642

49. Rothschild D, Susanto TT, Sui X, Spence JP, Rangan R, Genuth NR, Sinnott-Armstrong N, Wang X, Pritchard JK & Barna M (2024) Diversity of ribosomes at the level of rRNA variation associated with human health and disease. Cell Genomics 4: 100629

50. Saxena S, Cabuy E & Caroni P (2009) A role for motoneuron subtype–selective ER stress in disease manifestations of FALS mice. Nat Neurosci 12: 627–636

51. Sévigny M, Bourdeau Julien I, Venkatasubramani JP, Hui JB, Dutchak PA & Sephton CF (2020) FUS contributes to mTOR-dependent inhibition of translation. Journal of Biological Chemistry 295: 18459–18473

52. Shrestha P, Ayata P, Herrero-Vidal P, Longo F, Gastone A, LeDoux JE, Heintz N & Klann E (2020a) Cell-type-specific drug-inducible protein synthesis inhibition demonstrates that memory consolidation requires rapid neuronal translation. Nat Neurosci 23: 281–292

53. Shrestha P & Klann E (2022) Spatiotemporally resolved protein synthesis as a molecular framework for memory consolidation. Trends Neurosci 45: 297–311

54. Shrestha P, Shan Z, Mamcarz M, Ruiz KSA, Zerihoun AT, Juan C-Y, Herrero-Vidal PM, Pelletier J, Heintz N & Klann E (2020b) Amygdala inhibitory neurons as loci for translation in emotional memories. Nature 586: 407–411

55. Sidrauski C, Tsai JC, Kampmann M, Hearn BR, Vedantham P, Jaishankar P, Sokabe M, Mendez AS, Newton BW, Tang EL, et al (2015) Pharmacological dimerization and activation of the exchange factor eIF2B antagonizes the integrated stress response. Elife 4

56. Sinha NK, McKenney C, Yeow ZY, Li JJ, Nam KH, Yaron-Barir TM, Johnson JL, Huntsman EM, Cantley LC, Ordureau A, et al (2024) The ribotoxic stress response drives UV-mediated cell death. Cell 187: 3652–3670.e40

57. Sohn PD, Huang CT-L, Yan R, Fan L, Tracy TE, Camargo CM, Montgomery KM, Arhar T, Mok S-A, Freilich R, et al (2019) Pathogenic Tau Impairs Axon Initial Segment Plasticity and Excitability Homeostasis. Neuron 104: 458–470.e5

58. Storkebaum E, Rosenblum K & Sonenberg N (2023) Messenger RNA Translation Defects in Neurodegenerative Diseases. New England Journal of Medicine 388: 1015–1030

59. Strang KH, Golde TE & Giasson BI (2019) MAPT mutations, tauopathy, and mechanisms of neurodegeneration. Laboratory Investigation 99: 912–928

60. Suloh H, Ojha SK, Kartawy M, Hamoudi W, Tripathi MK, Bazbaz W, Schottlender N, Ashery U, Khaliulin I & Amal H (2025) Shared early molecular mechanisms revealed in P301S and 5xFAD Alzheimer’s disease mouse models. Transl Psychiatry 15: 97

61. Thelen MP & Kye MJ (2020) The Role of RNA Binding Proteins for Local mRNA Translation: Implications in Neurological Disorders. Front Mol Biosci 6

62. Tracy TE, Madero-Pérez J, Swaney DL, Chang TS, Moritz M, Konrad C, Ward ME, Stevenson E, Hüttenhain R, Kauwe G, et al (2022) Tau interactome maps synaptic and mitochondrial processes associated with neurodegeneration. Cell 185: 712–728.e14

63. Trinh MA, Kaphzan H, Wek RC, Pierre P, Cavener DR & Klann E (2012) Brain-Specific Disruption of the eIF2α Kinase PERK Decreases ATF4 Expression and Impairs Behavioral Flexibility. Cell Rep 1: 676–688

64. Vanderweyde T, Apicco DJ, Youmans-Kidder K, Ash PEA, Cook C, Lummertz da Rocha E, Jansen-West K, Frame AA, Citro A, Leszyk JD, et al (2016) Interaction of tau with the RNA-Binding Protein TIA1 Regulates tau Pathophysiology and Toxicity. Cell Rep 15: 1455–1466

65. Wang Y & Mandelkow E (2016) Tau in physiology and pathology. Nat Rev Neurosci 17: 22–35

66. Wolin SL & Walter P (1988) Ribosome pausing and stacking during translation of a eukaryotic mRNA. EMBO J 7: 3559–3569

67. Wu CCC, Peterson A, Zinshteyn B, Regot S & Green R (2020) Ribosome Collisions Trigger General Stress Responses to Regulate Cell Fate. Cell 182: 404–416.e14

68. Yoshiyama Y, Higuchi M, Zhang B, Huang S-M, Iwata N, Saido TC, Maeda J, Suhara T, Trojanowski JQ & Lee VMY (2007) Synapse Loss and Microglial Activation Precede Tangles in a P301S Tauopathy Mouse Model. Neuron 53: 337–351

69. Zhou C, Zhang M, Murray J, Paulo J, Gygi S, Shao S, Whitman M & Keller T (2025) GCN1 couples GCN2 to ribosomal state to initiate amino acid response pathway signaling. Science (1979) 390

70. Zyryanova AF, Kashiwagi K, Rato C, Harding HP, Crespillo-Casado A, Perera LA, Sakamoto A, Nishimoto M, Yonemochi M, Shirouzu M, et al (2021) ISRIB Blunts the Integrated Stress Response by Allosterically Antagonising the Inhibitory Effect of Phosphorylated eIF2 on eIF2B. Mol Cell 81: 88–103.e6

